# Context-dependent multiplexing by individual VTA dopamine neurons

**DOI:** 10.1101/408062

**Authors:** Kremer Yves, Flakowski Jérôme, Rohner Clément, Lüscher Christian

## Abstract

Dopamine (DA) neurons of the ventral tegmental area (VTA) track external cues and rewards to generate a reward prediction error (RPE) signal during Pavlovian conditioning. Here we explored how RPE is implemented for a self-paced, operant task in freely moving mice. The animal could trigger a reward-predicting cue by remaining in a specific location of an operant box for a brief time before moving to a spout for reward collection. *In vivo* single-unit recordings revealed phasic responses to the cue and reward in correct trials, while with failures the activity paused, reflecting positive and negative error signals of a reward prediction. In addition, a majority of VTA DA neurons also encoded parameters of the goal-directed action (e.g. movement velocity, acceleration, distance to goal and licking) by changes in tonic firing rate. Such multiplexing of individual neurons was only apparent while the mouse was engaged in the task. We conclude that a multiplexed internal representation during the task modulates VTA DA neuron activity, indicating a multimodal prediction error that shapes behavioral adaptation of a self-paced goal-directed action.

## Introduction

In the interest of survival, animals navigating through the environment adapt their behavior when reward contingencies change. Predicting a reward based on prior outcome contributes to the learning process for motor output through phasic response in midbrain DA neurons. Such RPE signaling promotes the formation of cue-reward associations ^1^. RPE signaling can be observed in Pavlovian conditioning ^2^ where the animal has little agency. This is particularly apparent for overtrained animals where the phasic response shifts from the reward to the cue during the formation of a novel cue-reward association. RPE coding scales with reward value ^3^, reward probability ^4^ and timing between events ^5^. If preceded by several stimuli, the signal shifts to the earliest reliable reward-predicting cue ^6^.

In naturalistic settings RPE computations may be more complex, possibly through the inclusion of hidden states inferred from previous training or subjective valuation ^7, 8^. For example, in self-paced operant behavior, the activity of midbrain DA neurons and the ensuing DA transients in the NAc may correlate with operant responding during drug or food seeking ^9, 10^. Additional studies suggest that DA neurons can also code for motivation (Hamid et al., 2016), action initiation ^11^, distance to reward ^13^ and time between stimuli ^14^. This raises the question, whether a single VTA DA neuron encodes multiple behavioral variables at the same time. It has been proposed that the phasic component of DA neuron firing encodes the external events (such as a cue and reward) while more sustained changes in tonic firing in the same cell would encode movement vigor or motivation ^15^. The simultaneous encoding of rewarding and motor variables in DA neurons is supported by recent findings in head fixed mice monitoring neural activity with calcium imaging in a virtual-reality environment ^16^. However, to the best of our knowledge, no study has taken advantage of the high resolution of electrophysiological recordings of action potentials to resolve tonic and phasic activity within individual neurons in freely moving mice executing a self-paced operant task.

To meet this challenge, we designed a spatial task by combining an operant behavior test originally developed to study hippocampal place cell firing patterns ^17, 18^ with food approach ^19^. In brief, mice had to find an unmarked “trigger zone” in the operant box and remain there two seconds, which would activate a light cue. Once the cue was presented, the animal had four seconds to collect the reward by licking for a drop of fat solution at a spout located at the other side of the box. The animal could engage in the next trial at its own pace. We performed single-unit recordings of the VTA and video recorded the movement of the mouse. We found that VTA DA neurons multiplexed phasic responses to salient events with tonic activity reflecting parameters of the motor output.

## Results

### Operant spatial task and behavioral performance

We injected a virus expressing cre-dependent channelrhodopsin (ChR2) and implanted a 16-channel optrode mounted into a microdrive into the VTA of DAT-Cre mice (Fig. 1a). After recovery, we started the pre-training phase that lasted five to ten days where the mice were conditioned in a cue-reward paradigm (Fig.1b,c). The mice learned to associate a randomly occurring 4s light stimulus (cue) with the availability of a drop of a fat solution (5% of lipofundin, BBraun, Sempach, Switzerland). We then switched to the cue-guided spatial navigational task for five to twenty days (Fig. 1c bottom timeline), where the cue was triggered once the mouse had spent 2s in a small (4×4cm), unmarked trigger zone (TZ) of the operant chamber (grey dotted square in Fig.1b). To collect the reward the mouse had to move to the other end of the box.

**Fig. 1.**
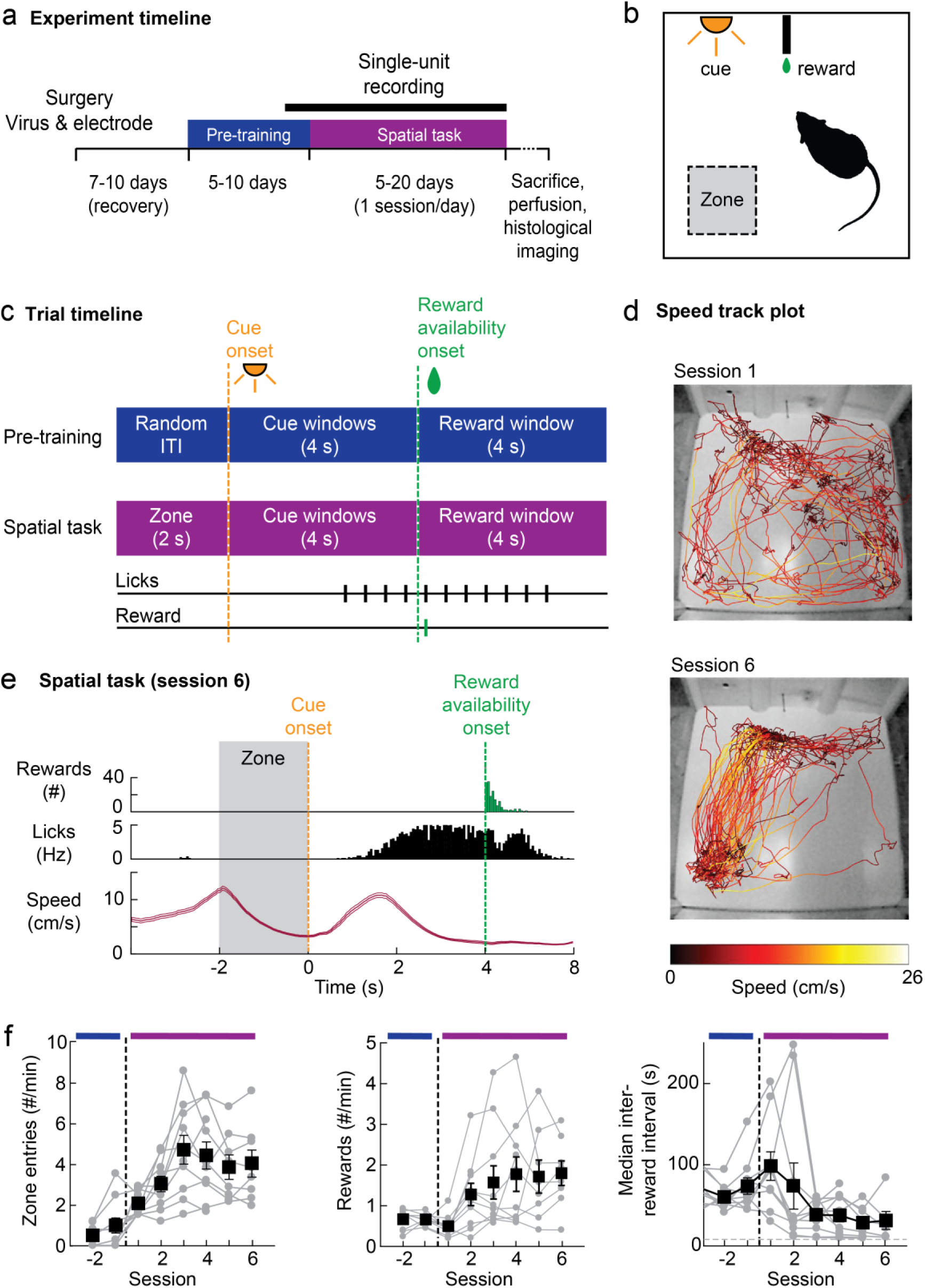
Operant spatial navigational task and behavioral performance. **a,** Experimental timeline. **b,** Schematics of goal-directed spatial task. The trigger zone is depicted here in grey but invisible for the mice that have to make the association between their spatial position and the cue to get a single drop of fat solution at the spout. **c,** During pre-training, mice have to associate a randomly occurring 4s light cue to the availability of the liquid reward. Once this association had been learned, we switch to the spatial operant task during which mice have to wait for 2s in the zone to trigger the light cue indicating that the reward can be collected. **d,** Track plots from 10.000 video frames (∼6-7 min) showing the locomotor pattern of the same mouse at the first and sixth session of the spatial task. **e,** Behavioral variables recorded during session 6 are shown in panel d: rewards collected during the 4s reward window (green), individual licks (black) and speed (dark red, mean ± s.e.m.) are aligned to the cue. f, Behavioral performance of all mice (n=10) for the last pre-training (−2 to −1) and the first spatial task (1 to 6) sessions. The black squares and error bars are the means ± s.e.m. across mice.

Within a few sessions, all mice found the TZ and the reward rate increased whereas the median inter-reward interval decreased in the first days (Fig. 1f and Fig. S1). Since the median inter-reward interval was insensitive to slow initiation or occasional breaks, we chose it as learning metric. The maximum number of reward was limited only by the duration of the executed sequence (approx. 10s). After three to five days, the mice executed the task using the shortest trajectories between the TZ and the drinking spout (Fig. 1d,e and Suppl. Video S1).

### In vivo recording and classification of VTA neurons

During the spatial task, we recorded the spike pattern of 159 neurons located within the VTA (Fig. 2a), which exhibited a wide range of firing patterns (Fig. 2b). We optogenetically tagged 16 cells as DA neurons, two of which were recorded simultaneously exhibiting contrasting responses to the cue and reward (Fig. 2d and S2c; see methods). One responded exclusively to the cue, whereas the other showed robust responses to cue and reward, which goes to show that with a given behavioral performance RPE coding may widely vary.

**Fig. 2.**
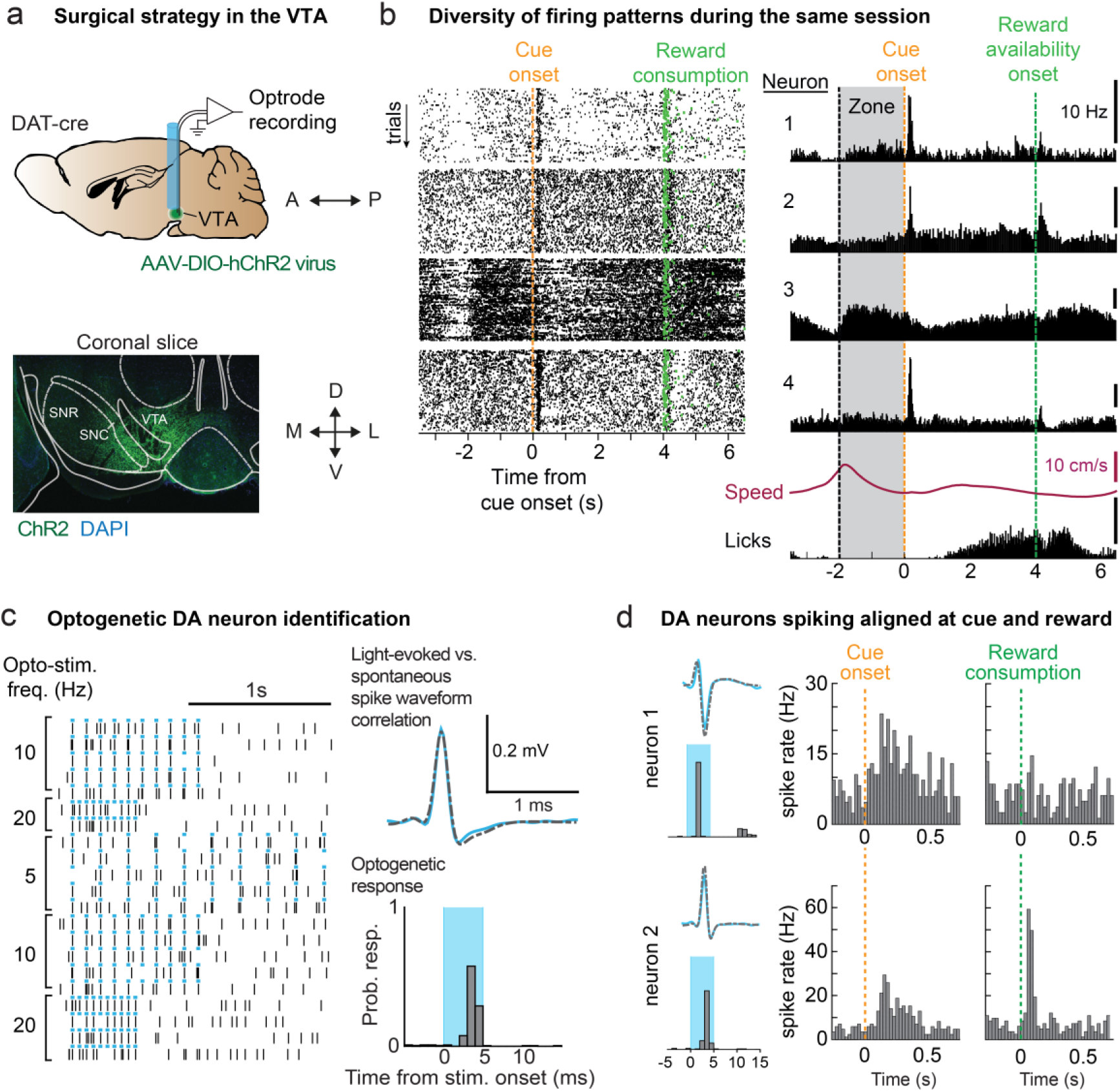
VTA *in vivo* recording and optogenetic tagging. **a,** Top: schematics of the single-unit recordings. Custom-made optrodes were implanted into the left VTA of adult DAT-cre mice that were injected with virus expressing floxed ChR2. Bottom: example of a coronal slice showing ChR2 expression (green) and DAPI staining (blue) in the VTA. **b,** Example of four simultaneously recorded units. Left: raster plots for each individual neuron. Right: average spiking histogram aligned to the cue showing the diversity of firing patterns during the same session along with the speed of the mouse and licking (below). Neurons 1, 2 and 4 were subsequently classified as DA neurons. **c,** Optogenetic identification of DA neurons. Left: 5ms blue light pulses at 5, 10 and 20 Hz. Neurons exhibiting a response rate ≥0.8 spikes/pulse during the 6 ms window after light onset and showing a waveform correlation ≥0.85 were identified as light-responsive (bin size opto=1ms). **d,** PSTHs of two DA neurons (bin size: 25 ms) which are light-responsive (neuron 1: opt.resp. = 0.819, waveform corr. = 0.952; neuron 2: opt.resp. = 0.8, waveform corr. = 0.996). They were recorded simultaneously during the spatial task at the same session but display different responses to external events. Both are aligned to the cue onset (left PSTH) and the reward consumption (right PSTH). The first neuron respond only to the cue while the second respond to both cue and reward (see raster plot in Fig.S2c).

We then aligned the cells to five events of interest (cue onset, reward consumption, lick, speed peak and acceleration peak, Fig.3a & S3a) and quantified the firing rate changes by computing the area under the receiver-operating characteristic (auROC) for each cell ^20^. A hierarchical clustering (Fig. 3a; see methods) yielded nine clusters, one of which (#5) contained 72 neurons whose most prominent features were large, phasic cue- and reward-evoked responses. All optogenetically identified DA neurons were assigned to cluster 5 and a comparison between light responsive and light non-response neurons failed to reveal any difference. (Fig. 3c).

**Fig. 3.**
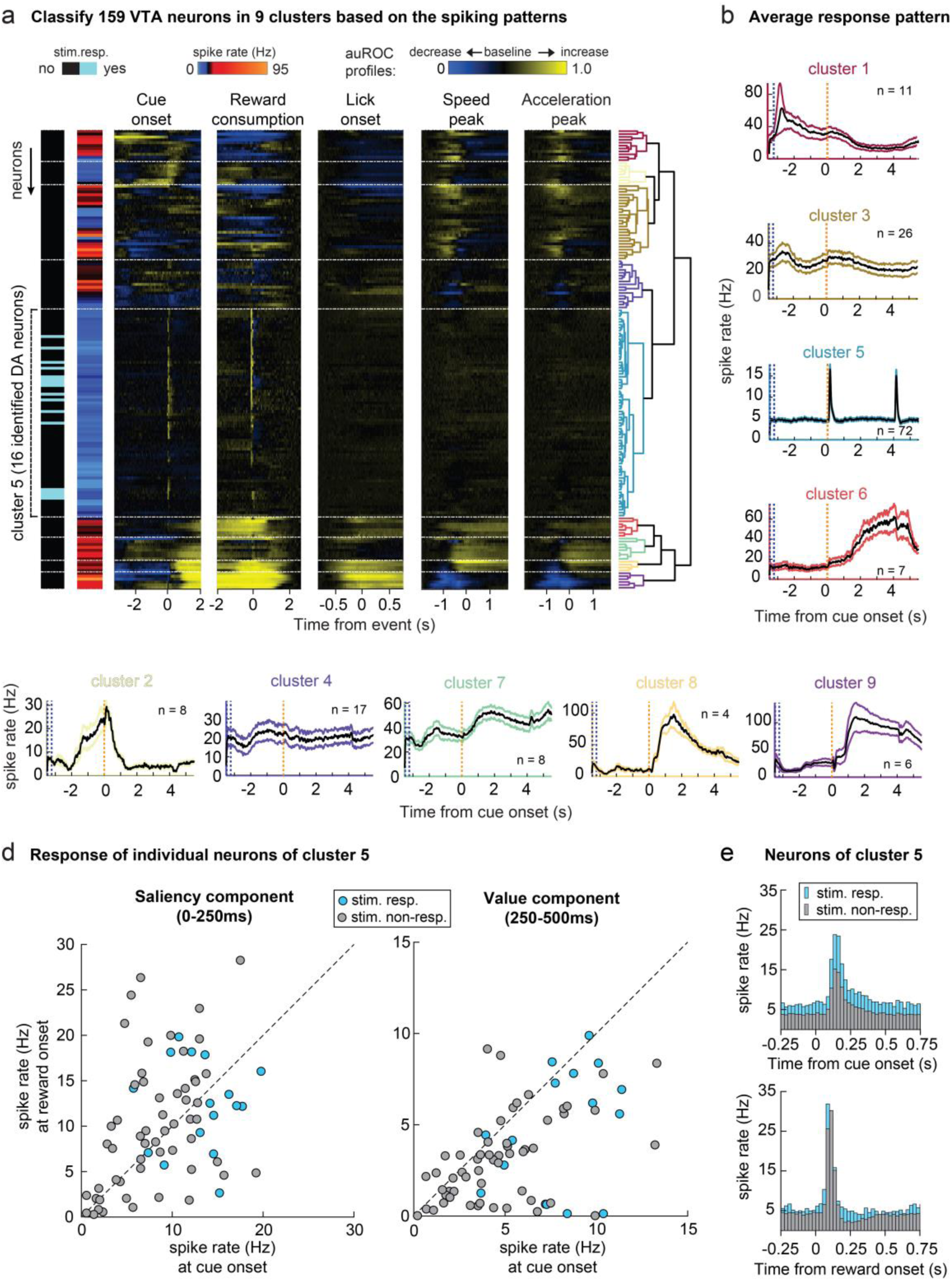
Clustering analysis. **a,** Hierarchical clustering of 159 VTA single-units (one per row, n=9 mice) based on their auROC spiking profiles (yellow: increase from baseline, blue: decrease from baseline), for five events and their mean firing rate. All 16 optogenetically identified neurons fell into cluster 5, which exhibits large phasic responses around the cue and reward. **b,** Average continuous spike density function aligned to the cue (mean ± s.e.m.) for functional cluster response patterns. Some clusters display task-related modulations, e.g. speed peaks (1 & 3), cue/reward cluster (5) or licking (6). Colors of the bounding boxes correspond to colors of the clusters in the dendrogram in panel (a). **c,** Average cue vs. reward responses during the saliency (0-250ms) and value (250-500ms) post-event window for all individual neurons of the cluster 5 (bin size PSTH=25ms). The spike rate between light responsive and light non-responsive neurons is undistinguishable. **e,** Average response histograms of all neurons of cluster 5, namely the optogenetically identified and putative DA neurons, to cue onset (left) and reward consumption (right) showing average responses to both events.

To start the spatial task, all mice had to complete the pre-task training and thus assimilated the cue-reward connection such that the reward was fully predictable. Nevertheless, in 32% (23/72 neurons), the response to the predicted reward was larger than the cue response, whereas 44% (32/72 neurons) showed the converse (Wilcoxon rank sum test, p<0.05), with only 16% (12/72 neurons) having no response at reward consumption. Again, the distribution of response amplitudes measured in optogenetically identified DA neurons (4/16 with [reward resp.] > [cue resp.] vs. 11/16 with [cue resp.] > [reward resp.]; Wilcoxon rank sum test, p<0.05), was similar to responses across all recorded DA neurons (Fig. 3d). Light non-response neurons of cluster 5 can thus be considered as putative DA neurons. The neurons of the other clusters showed sustained firing aligned to specific time points of the trial or were activated during specific motor output such as licking or locomotion. In line with previous studies ^8, 16, 20^, our analysis also revealed clusters reflecting motor output, which were modulated by locomotor speed or acceleration (cluster 1, 3 & 7; n=11, 26 & 8 neurons) or licking (cluster 6; n=7 neurons) as displayed in continuous spike density functions (Fig. 3b). Hereafter, we mainly focused on the DA neurons (cluster #5, n=72). We consider that the neurons of the other clusters, typically characterized by a high mean firing rate (>20Hz) and the absence of a clear phasic response at the cue and/or reward, are non-DA neurons (n=87 altogether).

### VTA DA neuronal pause during failed trials

In failed trials, during which the mouse did not wait for 2s in the TZ, no cue was presented. Yet, the mouse still approached the spout and started licking but did not get any reward (Fig. 4a-d). We monitored the neural activity along with the behavioral parameters distance, speed and licking for all DA neurons (n=72). We grouped neurons by animal and sorted them by behavioral performance measured as the median inter-reward interval (Fig. 4d, left). We found that phasic firing decreased in 23/56 (41%) putative DA neurons and 10/16 (63%) of optogenetically identified DA neurons, which manifested as a significant spike pause (Fig. 4d-f & Fig. S4). Discriminating neurons were not necessarily specific to a single event (Fig. 4g). VTA DA neurons can thus exhibit an internally generated error signal aligned to the mouse’s actions yielding a distinct activity pattern in the face of very similar motor output.

**Fig. 4.**
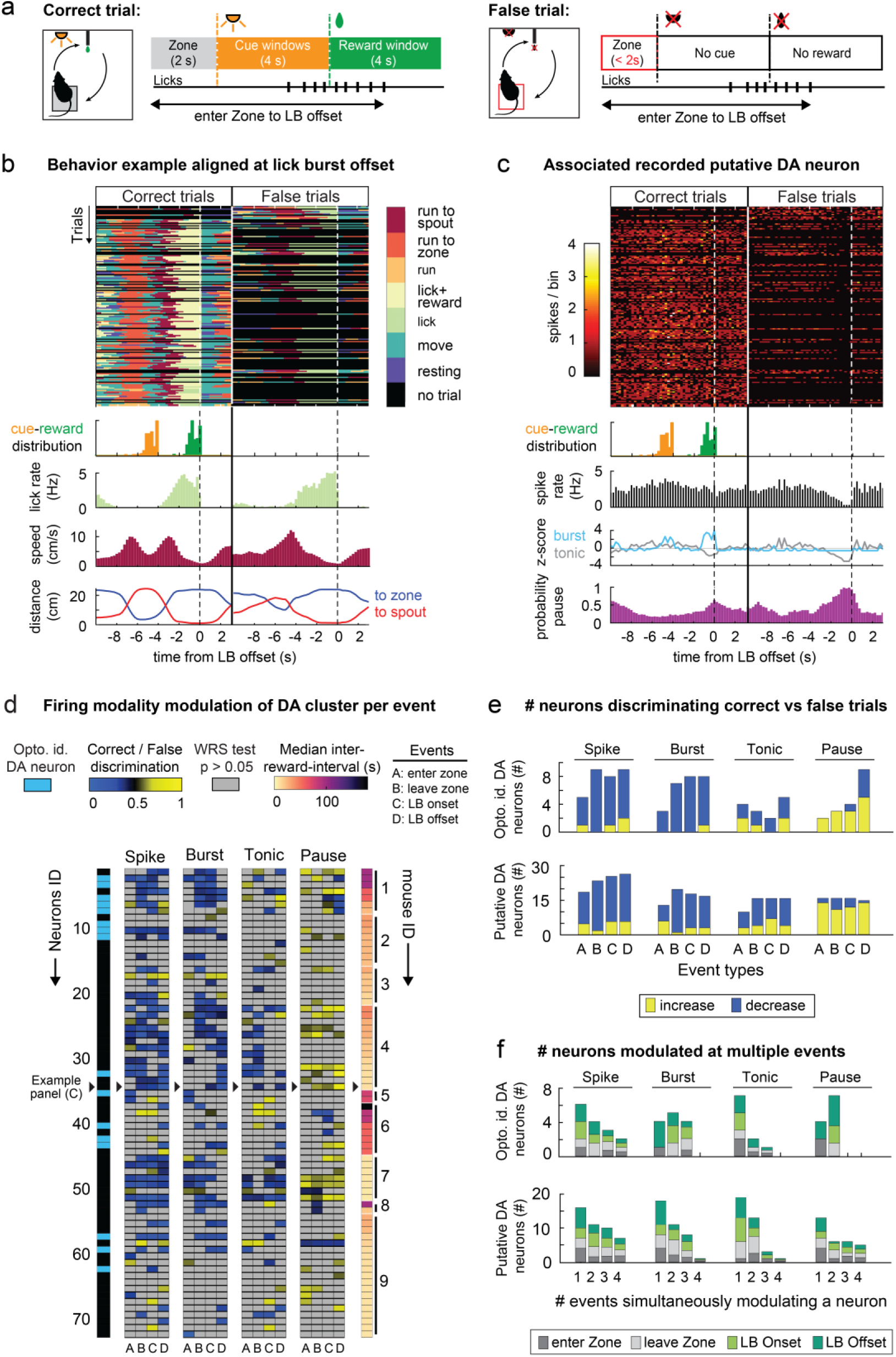
VTA DA neuronal pause during failed trials. **a,** Description of correct and false trials taking both place between the zone entry and lick burst (LB) offset. The animal failed to perform the task when it leaves the zone too early to trigger the cue. Note that only failed trials where the animal lick for the reward are considered. **b,** Correct versus failed trial analysis of one example session. The seven behavioral modules (run to spout, run to zone…) are defined in the methods. Trial-by-trial analysis of behavioral modules used to define correct (left) and false (right) trials across time. Below: cue (orange) and reward (green) distribution, lick rate (light green), speed (dark red) and the animal’s distance to the zone (blue) and to the spout (red) averaged across correct and false trials, respectively. **c,** Neuronal activity (bin size=250ms) of a putative DA neuron split into correct and false trials as defined from the behavioral analysis. Below: average spiking histogram (black), z-scored tonic (grey) & bursting (light blue) activity and probability of being in a spike pause (purple). Plots are aligned to the LB offset. **d,** Correct vs. failed discrimination of cluster 5 neurons (optogenetically identified and putative DA neurons, n=72). Significantly discriminating neurons are represented by their auROC change in spiking, burst firing, tonic firing and pausing. Black arrows highlight the example neuron in b. Neurons whose activity did not change significantly are represented in gray (Wilcoxon rank sum tests α=0.05). Neurons are grouped per animal and sorted by behavioral performance represented as the median inter-reward interval (see Fig. 1f; the shorter the interval, the better the animal’s performance). **e,** Number of neurons significantly discriminating correct vs. failed trials either with an increase or a decrease in the corresponding firing modality for optogenetically identified DA neurons (top) and putative DA neurons (bottom). **f,** The negative error signal, which manifests as a significant increase in pause probability for at least one event, is apparent in 62.5% of DA neurons (10/16) and by 41% of putative DA neurons (23/56). **g,** Number and distribution of events, which significantly modulate optogenetically identified DA neurons (top) and putative DA neurons (bottom).

### Neuronal modulation during goal-directed actions

We first analyzed the peri-event time histograms of each neuron around locomotion onset, time of maximal speed and lick onset within and outside the cue-reward window (Fig.5a). For example, alignment to the speed peak revealed a change in firing between the cue and reward onset but not when the alignment was made to the time of peak speed between trials (e.g. Fig.5b, bottom left). We computed significance in a 1s time window around the motor event ^21^ for all 159 VTA neurons (Fig. 5c and Suppl. Table 1). Up vs. down modulation was seen in both non-DA as well as DA neurons, whereby putative and identified DA neurons showed similar response profiles (Fig. S5 and Suppl. Table 1). For all DA neurons, we also examined the cue and reward responses and found that 53/72 showed such phasic responses. Importantly, 39/72 (54%) neurons encoded both, elements of motor behavior as well as cue-reward events. Such multiplexed modulation was more pronounced within the cue-reward window than outside (Fig. 5d). These results indicate that DA neuron modulation encodes motor action when it is goal-directed.

**Fig. 5.**
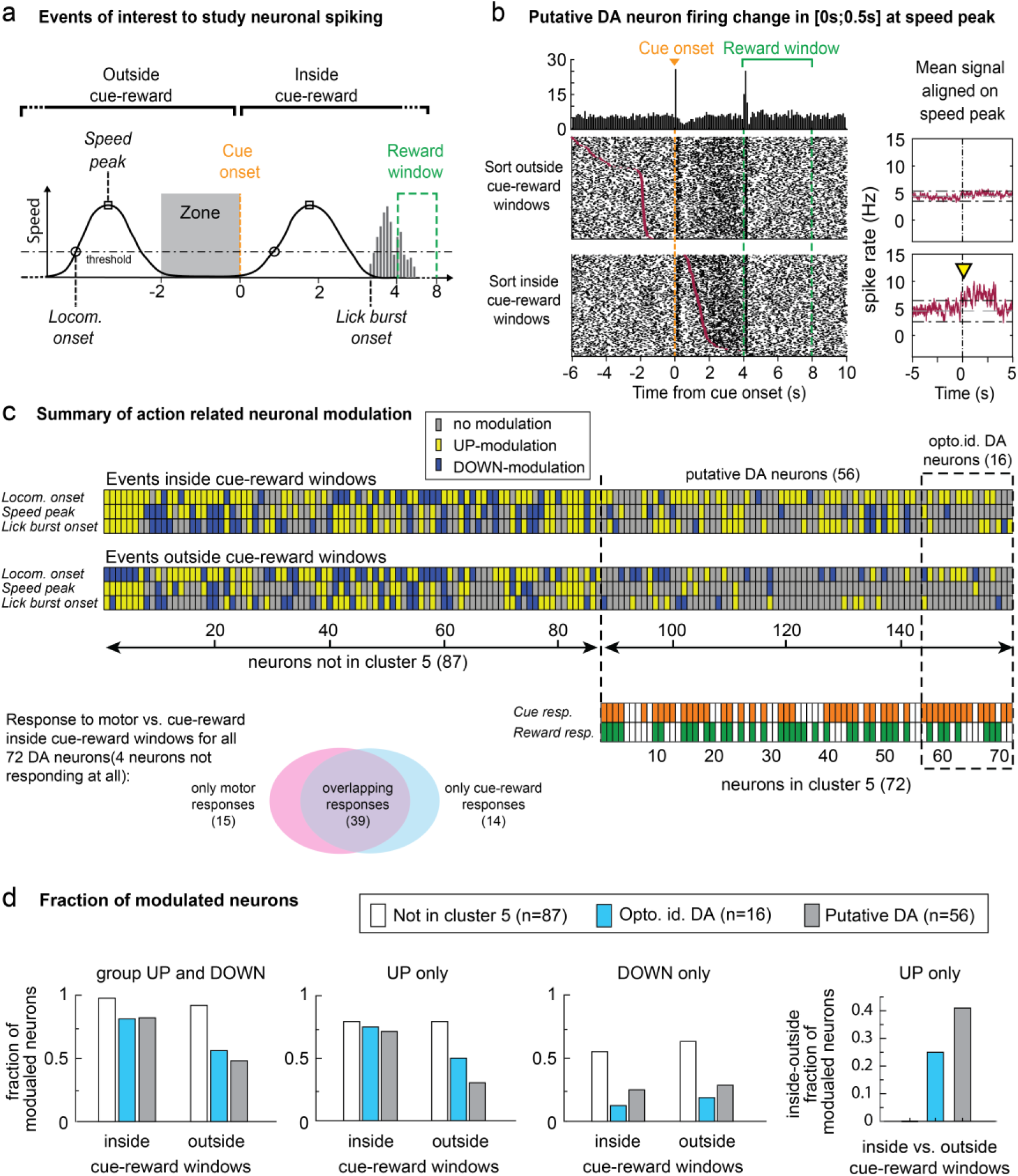
Neuronal modulation inside and outside of cue-reward interval. **a,** Schematics of three variables aligned at the trigger zone exit: locomotor onset, speed peak and lick burst onset. **b,** Left: PSTH (top) and raster plot (bottom) showing the firing of a putative DA neuron aligned to the speed peak (bin size = 100ms) inside and outside of cue-reward interval. Right: average spike rate aligned at the speed peak outside (top) and inside (bottom) of the cue-reward interval. The yellow arrow, which indicates the time of significant change of firing rate, is only observed when the trials are aligned to the speed peak inside of cue-reward interval. **c,** Display of significant modulations for all VTA neurons for the three variables. Bottom: Identification of phasic activity in all DA neurons and intersection between motor and cue-reward modulation (Venn diagram). Note that 39/72 neurons (54%) show modulation by both modalities, indicating multiplexing in a majority of DA neurons. **d,** The fraction of modulated neurons, depicted individually in panel c, grouped here by cell types and context. The UP-modulation is more frequent inside than outside of the cue-reward interval for DA neurons as shown on the most right panel.

### VTA DA neuron multiplexing in cue-reward intervals

To determine how many variables contribute to the spiking pattern of a single unit, we applied an encoding approach (Fig. 6a) for each neuron using a linear-nonlinear-Poisson (LNP) model, i.e. a generalized linear model (GLM) with an exponential nonlinearity and a Poisson noise commonly used to perform spikes encoding/decoding (see Methods). For a given session, we used seven variables (external events: zone, cue and reward as well as motor actions: lick, speed, acceleration and distance to the goal) and compared the resulting prediction to the recorded spiking through Pearson correlations (Fig. 6b). To compare the encoding during the task execution versus when the animal is not performing the task (Fig. 6c; see methods), we computed for each neuron the correlations for all correct trials’ time intervals (ranging from the zone entry to the lick burst offset) versus outside trials’ time intervals (excluding failed trials where the task is executed but wrongly). For DA neurons, the correlations was higher inside than outside of the task execution (Fig.6d-e). Note that the correlation is slightly larger when considering inside versus outside of the cue-reward intervals (Fig. S6a,b). This indicates that salient, reward-predicting events provide a time window during which goal-directed actions are closely tracked by DA neurons.

**Fig. 6.**
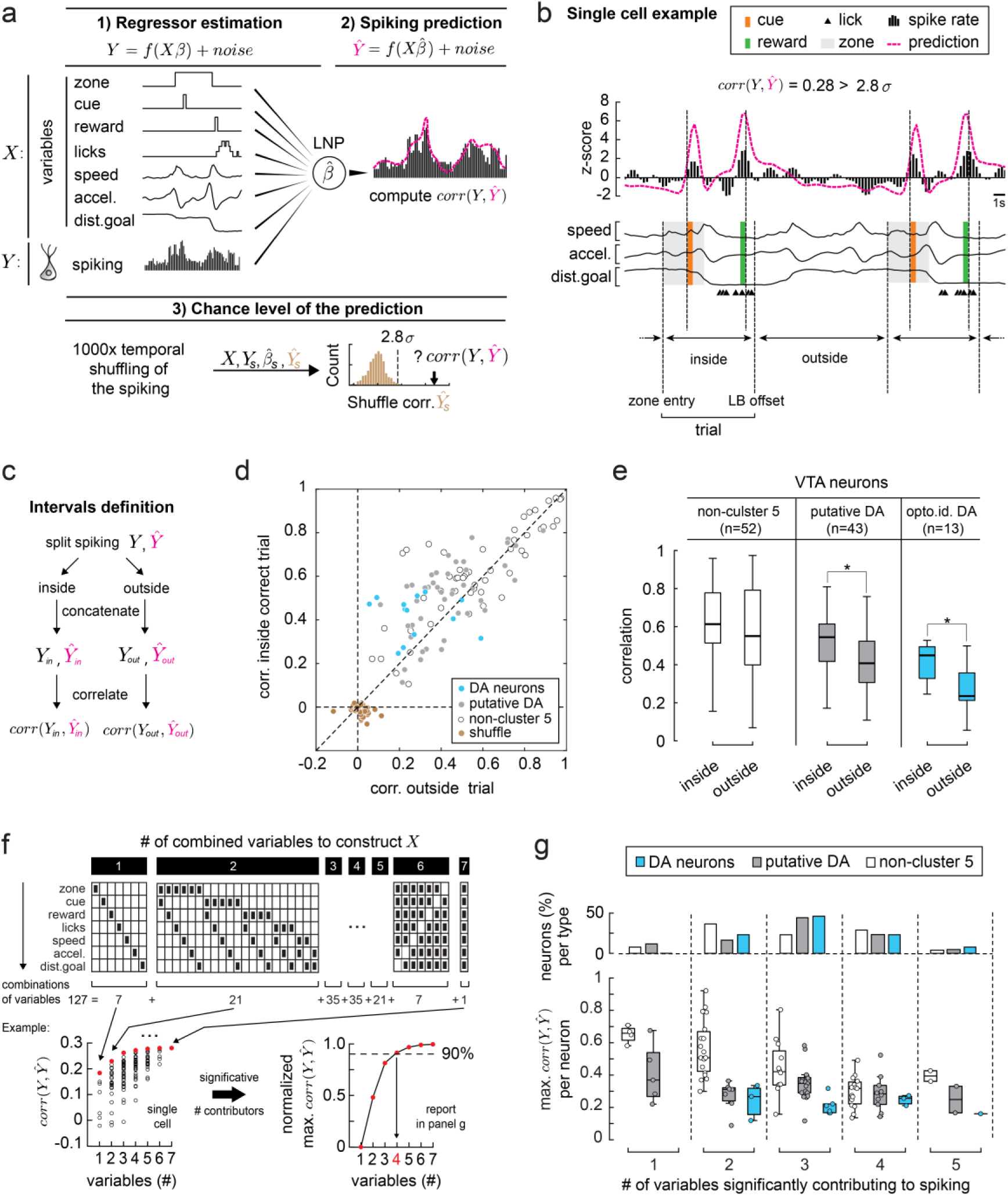
VTA DA neuronal multiplex encoding is specific to trial execution. **a,** Encoding model. For a given session, the design matrix X is constructed using seven behavioral covariates (events: zone, cue and reward as well as actions: lick, speed, acceleration and distance to the goal). Each activity vector Y is built from the recorded spiking (one per cell, 250ms bins). We first estimated the regressors ß for each neuron using a linear-nonlinear-Poisson (LNP) model, i.e. a GLM with an exponential nonlinearity f and a Poisson noise, for a sliding windows of 10 bins with lasso and elastic-net regularization to avoid overfitting. The prediction is compared to the true spiking through Pearson correlations. We proceeded only with neurons whose correlation is larger than chance level (n=118 → 108). **b,** Top: encoding example for a optogentically identified DA neuron. The true and predicted neural signals have been z-scored and displayed along with all behavioral variables. Bottom: in panels c, d and e we distinguish time periods inside of the trial (corresponding to the task execution taking place from the zone entry to the LB offset) and outside of it. **c,** We concatenate separately the recorded Y and predicted Ŷ spiking for inside the task execution on one hand and outside of the task execution on the other. We then computed the associated correlations. **d,** Scatterplot comparing the correlation of inside correct trials versus outside of the trials (excluding failed trials where the mouse executes the task but wrongly) as defined in b and c. **e,** Using a Kruskal-Wallis nonparametric one-way ANOVA with post-hoc test Tukey-Kramer, we show that the variables encoding is stronger during the task execution (zone entry to lick burst offset) than outside for all putative and optogenetically identified DA neurons (*: p<0.05). **f,** Illustration of the method to computed the number of variables significantly contributing to the spiking for each cell in panel g. Upper part: For a given cell, we performed the encoding analysis as depicted in panel a for each variables combinations (127 in total) constructing X by setting the missing variables to their mean value over time. Lower part: we selected the combination with the minimum number of variables necessary to cross 90% of the largest value among the computed maximum correlations (red dots); in the depicted example we get four significantly contributing variables (the arrow, in red). **g,** Lower part: shows maximum correlation for each neuron beyond chance level (n=108) as a function of the number of variables significantly contributing to the spiking following the method depicted in e. Upper part: percent of neuron per type and per number of significant behavioral variables; considering one or two variables, we have more non-cluster 5 neurons than identified and putative DA (44% vs 23% and 27%) while for three or more variable the tendency is opposite (56% vs. 77% and 72%). Moreover, the more a neuron encode variables, the lower the maximum correlation tend to be on average (0.53, 0.36, 0.33, 0.27 and 0.26 from one to five variables).

Finally, we estimated the number of variables significantly contributing to the spiking for each cell. In brief, we performed the encoding analysis for each variables’ combination (Fig. 6f top; see methods) by setting the missing variables to their mean value over time for each cell. The number of significantly contributing variables was obtained by selecting the combination with the minimum number of variables necessary to cross 90% of the largest value among the computed maximum correlations (Fig.6f bottom). We found that DA neurons typically required 3 or more contributors to explain their firing pattern based on both external events, i.e. cue and reward, and actions such as licking or speed. On the other hand, non-DA neurons typically encode 2 or less variables such as speed and acceleration (Fig. 6g). Finally, to determine the relative importance of the different variables for each group of studied neurons, we computed the fraction of correlation lost by removing one variable. As expected, external events (cue and reward) as well as licking were more important for DA neurons while speed and acceleration matter more in non DA neurons (examples in Fig. S6d and summary in Fig. S6e). Taken together LNP encoding confirmed that multiple variables determine the firing pattern of VTA neurons. More variables contributed in DA neurons than non DA neurons, and in the former the number was highest during the task execution.

## Discussion

Our single-unit recordings in freely moving mice show that during a navigational operant task, external events and motor related activity are encoded simultaneously in the majority of VTA DA neuron using distinct firing modes. The multiplexing was prominent during the task execution in the interval between cue presentation and reward collection.

A large fraction of neurons responded strongly even to a predicted reward, which is at odds with the classical RPE hypothesis ^6^. This cannot be due to different stages of learning, as all mice had fully associated the cue with the availability of the reward already during the pre-training well before the spatial task was introduced. Moreover, during sessions where several DA neurons were recorded simultaneously, we found some DA neurons showing large responses to the reward along with cue-responsive DA neurons that only showed a very small reward response. Moreover some DA neurons responded only weakly to the reward during the pre-training phase (Fig. S3), where the mice were still learning the cue-reward association (Fig.1c), ruling out the possibility that the introduction of the spatial component could have interfered with the RPE computation. A possible explanation for the persistent phasic response to the reward might be the variable time interval between cue and reward, which may degrade predictability. It is therefore likely that in self-paced tasks the phasic DA response to the reward never fully disappears.

Alternatively, the persistent phasic responses could represent an early saliency signal rather than a late value coding ^22^. Indeed, we found that the cue responses contained early saliency (0-250ms) as well as late value components (250-500ms), whereas reward responses predominantly occurred during the early 250ms window (Fig. 3d). Finally, it has been proposed that sensory prediction error is not represented by individual DA neurons but rather encoded in the firing pattern of many DA neurons ^23^. Thus, the diversity in responses observed here would simply reflect the necessity of a multidimensional encoding of prediction error to handle a self-paced complex navigational task.

In favor of RPE coding, we observed a robust decrease in spiking for failed trials, indicating that erroneous actions produced significant decreases in spiking causing sometime an actual pause. Previous studies have argued that VTA DA neuron pauses represent a negative RPE ^24, 25^, which in our case may not be limited to evaluation of the reward but also encompass actions in general. In our spatial task, the animal would build a model of the task structure and continuously improve future outcomes by adjusting their actions ^26, 27^. Another example for dopamine driving general action refinement may be found in songbirds. When presented with a distorted auditory feedback during song learning, birds produce a similar negative performance signal in VTA DA neurons ^28^.

While previous studies using FSCV have argued for task-related variables in addition to RPE coding in DA transients ^12, 13, 29, 30^ they fell short of identifying individual DA neurons that would encode multiplexed information with high temporal resolution. As mentioned above, the phasic component of DA neuron firing may encode cue and reward while sustained changes in tonic firing in the same cell could encode goal-oriented locomotion ^15^ providing efficient means to transmit complex information through a single channel. Frequency-division multiplexing has been proposed for DA neuromodulation on theoretical grounds ^15, 31–33^ but experimental evidence remains scarce.

There is also evidence for functionally specialized subpopulations, for example between midbrain DA neurons of the VTA and SNc. The later were observed to encode kinematics regardless of the valence of trigger of the movement (Barter et al., 2015). Similarly monitoring the DA terminals in ventral and dorsal striatum revealed functional specialization ^34, 35^. In VTA, some DA neurons predominantly encoded reward-while other mainly movements or both ^36, 37^. One study reported burst at the onset and offset of voluntary movement in VTA DA neurons (Wang & Tsien, 2011). Finally, when monitoring the activity of many neurons with calcium imaging in head fixed mice navigating through a virtual-reality environment spatially organized subpopulations of DA neurons were identified that may encode information about a subset of behavioral variables in addition to encoding reward ^16^.

Using peri-event analysis ^21^ and a generalized linear model approach ^38^, we found a mixture of external events and goal-directed actions predicting the neural signal of VTA DA neurons (cluster 5 in Fig.3a and Fig.6 and S6) consistent with task-related modulation in DA firing rates described in monkeys while performing a forelimb reaching task ^39^. This was in contrast to non DA VTA neurons whose firing patterns could largely be explained by motor output. The fact that motor output alone did not reliably modulate the spiking of DA neurons but rather tracked specific actions during relevant time windows such as the cue-reward time-interval (or more generally the trial time interval ranging from the zone entry to the lick burst offset), suggests that it represents a learned signal. This is in line with optogenetic manipulations where direct stimulation of VTA GABAergic neurons show disruption of motor output ^40^, whereas phasic firing of VTA DA neurons reinforces action rather than driving motor output ^12, 41^. In SNc DA neurons, transient increases in firing have been observed before action initiation and optogenetic stimulation promoted movement initiation ^21^.

Multiplexing may arise from the convergence of many motor inputs onto VTA DA neurons, namely inputs from motor control (M1, M2) and general locomotor regions (e.g. diagonal band of Broca, medial septal nucleus, pedunculopontine nucleus and laterodorsal tegmentum) ^42–44^ and inputs shaping the RPE computation. The latter are believed to be more widely distributed, neurons contributing to the positive RPE signal have been found in striatum, ventral pallidum, subthalamic nucleus, pedunculopontine nucleus and lateral hypothalamus ^45^. For the negative error signals, the lateral habenula has emerged as the main input ^46, 47^. Interestingly specific inputs can differentially affect individual firing modes in DA neurons thereby precisely addressing a single channel of the multiplexed signal ^48^. Without surprise, optogenetic manipulations of VTA DA neurons have revealed strong behavioral effects, which may be explained by the many information streams treated in parallel ^43, 49–52^.

But how might the signal then be de-multiplexed? Downstream of the VTA, cell-type specific changes in NAc can occur during learning ^53^. In fact D1- and D2-receptors expressed on individual medium spiny neurons differentially regulate synaptic plasticity of afferent glutamate transmission ^54, 55^, which has been proposed to underlie reinforcement learning ^56^. As D1- and D2-receptors have different affinities, it makes them ideal candidates to decode a multiplexed DA-signal ^57, 58^, and differentiate between the phasic and tonic spiking. In line with this interpretation does closed-loop stimulation of either D1- or D2-striatal medium spiny neurons (MSNs) during movement execution bi-directionally shift specific parameters of the stimulated movement ^59^. Similar mechanisms could underlie the optimization of task-related parameters in ventral striatum during goal-directed actions. Similar DA-dependent mechanisms have been reported in other downstream areas of the VTA: stimulation of VTA DA projections to prefrontal cortex showed opposing effects between phasic and tonic stimulation ^60^ and functional imaging studies in humans have implicated the posterior medial frontal cortex in learning from errors and showed that it was dopamine and D2-receptor dependent ^61^.

We provide here experimental evidence for goal-directed multiplexing in individual DA neurons for freely moving mice. Cue-reward signals and movement related firing are processed in parallel thanks to the phasic and tonic modes of DA neurons. Each single unit is modulated in an outcome-dependent manner, thus contributing to the learning of new contingencies. This is a crucial step to understand how midbrain DA neurons work in natural settings. Our results may also have translational implications. Presentation of salient cues to patients with Parkinson’s disease can improve symptoms, for example by overcoming freezing of gait ^62, 63^. If elevated DA levels constitute an underlying mechanism, then our data indicate that adding cues may recruit otherwise silent VTA DA neurons (some of which are correlated with motor actions), which are less affected by neurodegeneration than SNc neurons ^64^ and could partially compensate in regions where SNc and VTA projections overlap. Another implication might be that in substance abuse, where pharmacological jamming of the error signal by the drug in addition to environmental cues could drive the persistent consumption despite negative consequences and lead to addiction ^65, 66^.

## Acknowledgments

We thank Anthony Holtmaat, Alexandre Pouget, Rui Costa, Wolfram Schultz, Léon Fodoulian, Michael Loureiro and Ruud van Zessen for helpful comments on the manuscript and all members of the Lüscher lab for stimulating discussions. This work was supported by the Swiss National Science Foundation (SNSF; FNS310030B_170266) and by the European Research Council (ERC; UE7-MESSI-322541).

## Author contributions

C.L and Y.K. designed the study and were responsible for project administration. C.L. was responsible for the funding acquisition. Y.K. and C.R. did the investigation. Y.K. was responsible for the project supervision. Y.K and C.R. did the virus injection, optrode implantation, electrophysiological and behavior recording, Y.K. did post-mortem histology and imaging, Y.K., C.R and J.F did the data curation, formal analysis and provided custom-written code. Y.K and J.F elaborated the methodology and data visualization. J.F. elaborated the encoding scheme. C.L., Y.K. and J.F. wrote and edited the manuscript with the input of all others.

## Key resources table

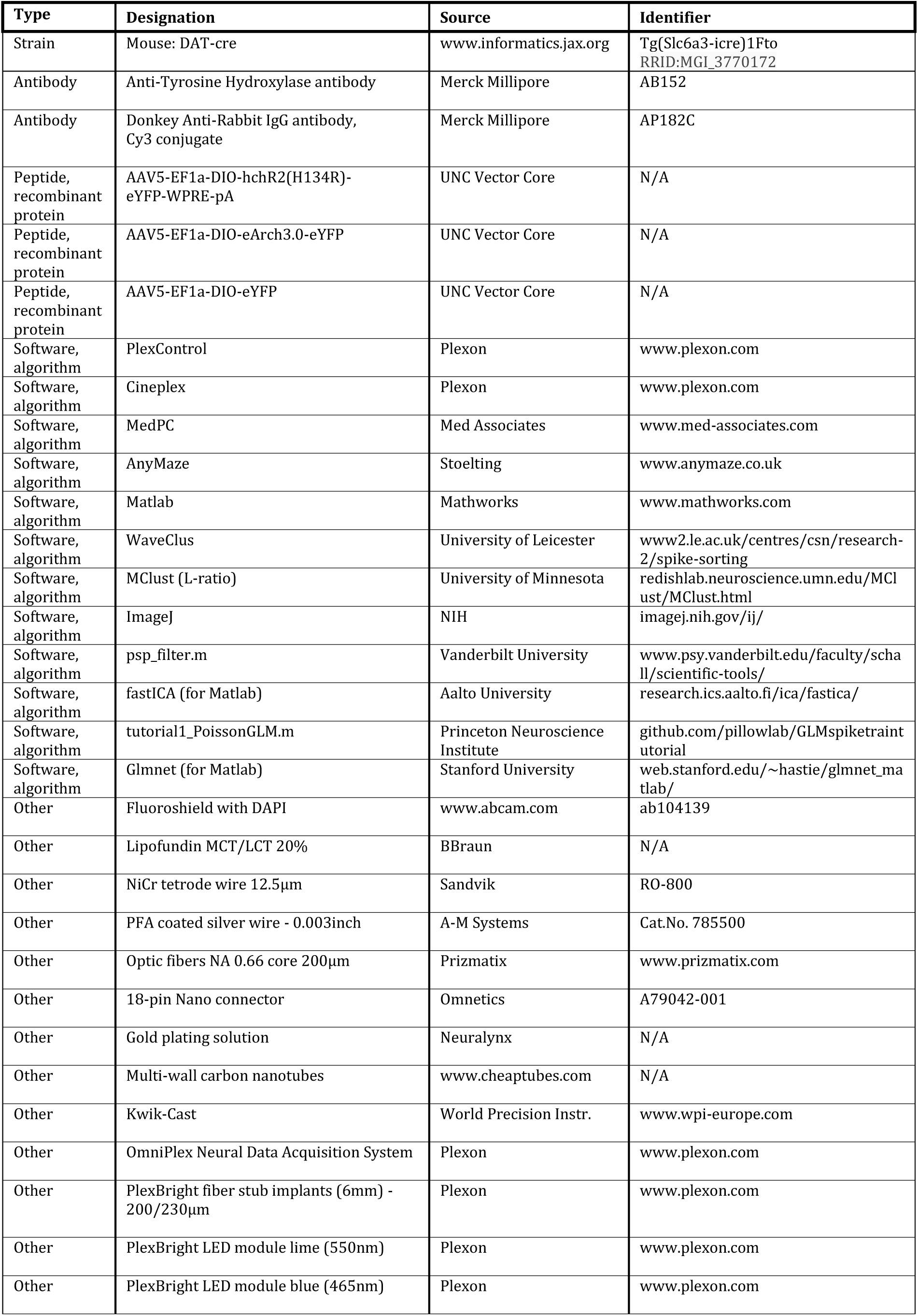

## METHODS

### Animals

All experiments were reviewed by the institutional ethics committee and approved by the relevant authorities of the Canton of Geneva. Animals were housed individually after implantation to avoid damage to the optrode. During training, they were food-restricted to a single pellet of chow per day (∼2g). The animal’s weight did not drop below 85%. All animals used in this study were adult DAT-cre mice. Only males were used for the electrophysiological recordings to minimize the impact of the weight of the optrode implant.

### Surgical procedures and viral injections

For recordings, adult male DAT-cre mice (n=10) were implanted with a custom-built 16-channel optrode mounted into a microdrive ^67^. 16 NiCr-microwires were assembled into an electrode bundle, glued to a 250 µm large NA (0.66) optic fiber (Prizmatix, London, UK) and mounted into the microdrive assembly. The tips were electroplated in gold solution containing multi-wall carbon nanotubes (1mg/ml)^68^ to reach a resistance of 100-150 kΩ using the NanoZ impedance tester. The completed implant did not exceed 2g in weight. During the surgery, mice were anesthetized using isoflurane (∼2%), the scalp was injected with lidocaine and the skull exposed. Three skull screws were inserted and a craniotomy performed above the VTA. The dura was carefully removed and 400-600nl of cre-dependent ChR2 were slowly injected into the VTA (AP: −3.2 ± 0.2; ML: 0.8 ± 0.2; DV: −4.2). Then, the optrode was lowered slowly into the brain using a micromanipulator and implanted just above the VTA (AP: −3.2 ± 0.2; ML: 0.8 ± 0.2; DV: −3.8 to −4.0). The space between the brain surface and the implant was first sealed using silicone elastomer (Kwik-Cast, WPI, Friedberg, Germany), then secured using super glue and dental cement. The ground electrode was inserted into the posterior part of the contralateral hemisphere devoid of brain vessels.

### Behavioral training

After the surgery, animals were given at least one week to recover. For recording experiments, DAT-cre mice were progressively food-restricted down to 1 pellet of chow per day to reach approximately 85-90% of their initial weight. During one habituation session in the behavioral apparatus (MedAssociates, Fairfax, VT) they located the drinking spout and had free access to the fat solution (solution of lipofundin 5% in water). Then mice started pre-training to associate a light cue with the availability of a liquid reward. If during the reward window mice licked against a custom-made piezo-based lickometer, they obtained a single drop of liquid reward. The analog piezo lickometer signal was sent to an Arduino board which applied a threshold and simultaneously sent a digital signal to the MedAssociates software and to our electrophysiology recording system for high temporal resolution timestamping of lick detection (Plexon). Lick bursts were defined as starting when the inter-lick interval (ILI) dropped below 0.5s and ended once the ILI exceeded 2s. The duration of the cue was 4s and the reward window lasted 4s. Both remained unchanged across all training stages. During pre-training, the average random inter-trial interval between two consecutive cues was increased across pre-training sessions from an average of 45s to 65s during the last session. Animals underwent pre-training until they exhibited good success rates (>0.8) of reward per cue while simultaneously reducing their licking between subsequent cues. During both, the pre-training and the spatial task, AnyMaze (Stoelting, Dublin, Ireland) or CinePlex (Plexon), were used to online video track the animal’s position (defined by its center of gravity). A 4×4 cm zone was defined inside the 22×22 cm operant chamber. Digital signals were sent to MedPC (MedAssociates) to trigger the light cue and reward delivery, if the mouse spent 2s inside the trigger zone as required by the task. Additionally, all zone entries, licks, cues and rewards were recorded with high temporal resolution by our electrophysiology recording system.

### Electrophysiological recordings and spike sorting

We started recordings during the last pre-training sessions when mice reached the criterion to move to the spatial task. If a neuron exhibited DA-like firing patterns or responded to the opto-identification protocol, the mouse was switched to the spatial task on the next day. Neuronal activity was recorded at 40 MHz sampling rate and all behavioral variables timestamped by our recording system (Plexon). To allow for unrestrained locomotion and to reduce weight, we used a dual LED and 16-channel commutator (Plexon). The distance between the headstage and the commutator was optimized for each animal. Recording sessions lasted 60-90 minutes, simple online sorting was performed to monitor recording quality. For analysis, offline spike sorting was performed using WaveClus ^69^. A frequency of 150Hz was used to separate the wideband signal into LFP and spike bands. Positive or negative thresholds were applied depending on the observed waveforms. Spike sorting results were visually inspected, the optimal temperature parameter in WaveClus selected and minimal manual correction applied. Only units with a L-ratio<0.05 were further analyzed ^70^. After each session, the microdrive was used to lower the optrode by ∼50µm and the same procedure applied on the next day. Once the electrode had crossed the VTA along the dorso-ventral direction, animals were killed and perfused to recover the location of the electrode track (Fig S2C). Cell-type specific expression of ChR2 was also verified (Fig S2B).

### Optogenetic identification of single-units

After each recording session and before the optrode was lowered for the next day, we delivered blocks of 10 pulses of 5ms using a LED (Plexbright LED modules, Plexon driven by a Master-8 stimulator, A.M.P.I., Jerusalem, Israel) at frequencies of 1, 2, 5, 10 and 20 Hz. Light intensities ranged from 1-5mW. Light evoked spikes were detected during a 6ms window after light onset. A single-unit was validated as an optogenetically identified DA neuron if its response rate was ≥0.8 and the correlation between the average light-evoked waveform and the average of all other spike waveforms produced during the entire recording session was ≥0.85.

### Post-mortem histology & imaging

After the optrode had crossed the VTA, mice were sacrificed with a lethal injection of pentobarbital (150mg/kg) and perfused with 4% of paraformaldehyde in cold PBS. After fixation, the optrode was slowly retracted using the microdrive and the brain extracted. Coronal brain slices of the VTA were cut 80-120-µm thick. To quantify the specificity of ChR2-expression, slices were immunostained for tyrosine hydroxylase (TH). Briefly, slices were permeabilized with 0.1% Triton in TBS-Tween 20 (TBST), blocked using 1% BSA in TBST and incubated with primary anti-TH antibody (dilution 1:500; AB152, Merck Millipore, Schaffhausen, Switzerland). 24h later, slices were rinsed and incubated with secondary antibody (dilution 1:500; AP182C, Merck Millipore) diluted in 1% BSA TBST for 1-2h. After each step, samples were washed with TBST.

Slices were mounted and covered on microscope slides using Fluoroshield mounting medium with DAPI (Abcam, Cambridge, UK), low-resolution imaging was performed using either a 5x/NA 0.25 or 10x/NA 0.45 objective in a slide scanner (Mirax or AxioScan Z1; Zeiss, Jena, Germany). High-resolution stacks were acquired in a confocal fluorescence microscope (LSM800 using 40x/NA1.4, Zeiss or A1r Spectral 40x/NA1.3, Nikon, Amsterdam, Netherlands).

In recorded animals, the electrode track could easily be located in 1-3 brain slices and was mapped to the stereotaxic coordinates of the slice where the track was most prominent. For cell counting, confocal high-resolution imaging stacks (z-steps of 2.5µm) were acquired, then cells expressing ChR2 and those stained for TH were counted manually using ImageJ.

### Event-related neuronal responses

For each neuron, spikes were binned into a regular time grid of 250ms time bins. Neurons with a mean firing rate <1Hz as well as maximal firing rate <3Hz were excluded leaving us with 215 neurons. For each neuron, we constructed five event-related profiles around cue onset, reward consumption, the mouse’s lick, speed peak and acceleration peak. The increase or decrease of each time bin from the baseline (defined for each event as [-3.5;-3.25] for the cue onset, [2.5;3.0] for reward consumption, [-1.0;-0.75] for lick onset, [-2.0;-1.75] for the speed and the acceleration peaks) was quantified using an area under a receiver-operating characteristic curve (auROC) method ^20^. auROC values are between 0 and 1, values <0.5 indicate a decrease relative to the baseline, while values >0.5 reflect increases in firing relative to the baseline. To visualize the absolute neuronal firing rate, for each neuron we also computed the corresponding continuous spike density function (cSDF) by convolving the spikes with an excitatory post-synaptic potential (ePSP) kernel ^71^ using the algorithm proposed by the Schall Lab ^72^. All analyses in this work were performed using custom written code in Matlab (Mathworks, Natick, MA).

### Clustering of neuronal firing patterns

We recorded a total of 215 neurons from 10 mice. We then applied a two-step clustering strategy to isolate neurons within the VTA. First, we applied a dimensional reduction to the five set of auROC profiles (i.e. cue, reward, lick, speed, acceleration) using an independent component analysis (ICA) (fastICA Matlab package). We then applied a hierarchical clustering, using the standardized Euclidean distance metric and ward linkage method, to the 18 combined ICA components plus the mean firing rate (i.e. a 19 x 215 matrix). At this stage, the information obtained from the optogenetic identification was not considered. The clustering quality was assessed with the mean silhouette index and cophenetic correlation coefficient. We identified the putative dopamine (DA) cluster by the typical phasic responses to cue and reward, which was confirmed by the fact that all optogenetically identified DA neurons (n=16) fell into this cluster (cluster #8 in Fig. S3a). By moving our optrode across the VTA, we further refined our set of neurons by accepting only single-units located between the first and the last optogenetically tagged DA neuron in a given animal. This resulted in 159 VTA neurons, which were subjected to a clustering procedure yielding 9 clusters (Fig. 3a).

### Definition of DA neuron firing modes

Since in the freely moving condition, we found elevated firing rates compared to head-fixed animals, we could not directly apply the 80/160ms rule ^73^ to separate spike bursts from tonic firing. We adapted the 80/160 rule by first calculating the mean inter-spike interval (ISI) across the entire session. The onset was then defined by the first interval shorter than a 1/3 of the mean ISI and the offset was detected by using an interval twice as large as the onset threshold. This resulted in burst detection thresholds similar to previous studies when DA neurons fired at 4-5Hz, but scaled for neurons with increased firing rates. All spikes not attributed to a spike burst were tagged as tonic spikes. To detect spike pauses, we plotted the histogram of ISIs of all tonic spikes (ISItonic). We then fitted a Gaussian in the interval [0;3] of this distribution and computed the time at which the difference between the fit and the actual distribution was largest. We used this value as the threshold for pause detection. If this threshold was smaller than 2, it was set to 2. In subsequent analyses, time bins were considered to be in a pause if they were between the spike starting a pause and the spike ending a pause.

### Behavioral modules

To characterize more precisely behaviors during each session, we defined seven modules based on the recorded behavioral variables: at rest, (slow) move, run, licks, rewarded licks, run toward zone and run toward reward. Transition points between resting and moving periods used a threshold of 2.1cm/s. We then computed the mean speed value for move intervals and used a move-run threshold of 2.8 cm/s thereby defining the resting, moving and running modules. Lick bursts were detected as described above, starting with a ILI ≤ 0.5s (lick burst onset) and ending when ILI ≥ 2s (lick burst offset). We defined rewarded lick bursts as those whose onsets were occurring between a cue and a reward. All other lick bursts were considered as unrewarded lick bursts. Finally, we computed the distance to the reward and to the zone in order to further specify if run modules were directed towards the zone or towards the reward.

### Comparing neural signal underlying correct and false trials

Correct trials lasted approximately 10s: the mouse entered the zone, waited for 2s in the zone until the cue turned on and finally ran toward the spout (or valve) where the reward was delivered from 4 to 8s after the cue. The mouse spent a few seconds near the spout to consume the reward and then returned to the zone. We sought to detect failed trials that had motor output which was similar to correct trials but during which neither the cue nor reward were delivered. For example, mice frequently left the zone before the CS came on, ran toward the spout, licked but did not receive the reward. Using the behavioral modules defined above, we defined false trials with similar speeds and trajectories to correct trials. In a failed trial, we required the time from the module “run toward the zone” to the module “run toward the valve” to last at least 3s to avoid the case when mice just randomly crossed the zone and ran toward the valve. To be included in failed trials, mice had to lick and test for reward delivery. Similarly, we used these same behavioral modules to identify correct trials and subsequently validated the procedure using the cue- and a reward-events, showing that this procedure was capable of extracting correct and false trials based on similar motor output.

Using the definition of successful and failed trials, we then compared their associated neural signal (spike, tonic, burst spiking and pause) around 4 events of interest (zone entry, zone exit, lick burst onset and lick burst offset). We applied an auROC analysis ^74^ to compare the mean signal in false trials to activity in correct trials for the four events. The response windows for the different events were: zone entry [-0.75;1.5], zone exit [-0.5;1], lick burst onset [0;1] and lick burst offset [-0.75;0.25]). To determine if the relative change was significant, we performed a Wilcoxon two-sided rank sum test (α=0.05). For cluster 5 neurons (identified and putative DA neurons having an average firing rate of 4-14Hz), the analyses were carried out on all spikes, tonic firing, burst spiking and pauses. In contrast, non-cluster 5 neurons were analyzed only with all spikes due to too larger average firing rates forbidding to use tonic firing, burst spiking and pauses as defined beforehand.

### Action related modulation of neuronal spiking

To determine which neurons were significantly modulated by goal-directed actions (licking and locomotion), we analyzed peri-event time histograms around lick burst onsets, locomotion onsets and speed peaks ^21^. In a window of [-5;5] seconds around these time points, spikes were time averaged in a sliding window of 100ms shifted by 1ms steps. The baseline was defined as the [-5;-4] time interval. If the neuronal activity during the response window ([-0.5;0.5] for locomotion onset, [0;0.5] for speed peak and lick burst onset) exceeded 3 standard deviations and remained above for at least 50ms, the neuron was considered significantly modulated by the action. When a minimum time interval was imposed between external events (cue and reward) and action onsets, we first removed all movement-related time points closer than 750ms to an external event and the same analysis was applied to the remaining behavioral events.

### Encoding analysis of the spiking

To determine which events and behaviors contributed most to the spiking output, we applied a Generalized Linear Model (GLM) to predict spiking using a Linear-Nonlinear-Poisson (LNP) model ^75^. The standard LNP model *Y* = *f*(βX) + ε is simply a GLM with an exponential nonlinear link function *f* and a Poisson noise ∊ commonly used to predict spiking ^76, 77^.

To improve between session comparison, this analysis was restricted to a homogeneous subset of 36 sessions (118 out of the 159 neurons). The single-units were selected based on the session duration (65.9 ± 3.52. minutes, mean ± s.e.m), i.e. only session with at least 30 minutes of continuous recordings were selected. The encoding procedure is depicted in Fig. 6a. Each activity vector *Y* is built from the recorded spiking (one per cell, 250ms bins). The design matrix X contained seven covariates (i.e. variables): three events (cue, valve and zone) and four behaviors (licks, speed, acceleration and distance to goal) resampled using same bin size as for the spiking. The presence inside the trigger zone was represented in a binary vector as were cue and valve events, indicated by binary events lasting 2 bins. The remaining continuous variables (licks, speed, acceleration and distance to reward) were normalized and z-scored. We used a sliding window of 10 bins to predict the spiking at a given time bin. We estimated the regressors β for each neuron using the LNP model. To avoid over-fitting due to the large number of parameters used to predict the spiking, we resorted to a Lasso and elastic-net regularization (glmnet for Matlab). Pearson’s correlation coefficient was used to compare the recorded and predicted spiking (e.g. Fig. 6b). Finally, we temporally shuffled the neural signal 1000x by a randomized time offset (50-500s) and performed the analysis again to check that the prediction is better than chance level (threshold of 2.807 standard deviations to the shuffled distribution). Accordingly, only neurons whose Pearson correlation was higher than shuffled one were used in the main figure (108 out of the 118 neurons).

To compare the encoding inside versus outside of the task execution time intervals, we concatenate separately the recorded Y and predicted Ŷ spiking of each neuron for inside the task execution on one hand and outside of the task execution on the other (Fig. 6c). We then computed the correlations for all correct trials’ time intervals (ranging from the zone entry to the lick burst offset) versus outside of correct + failed trials’ time intervals (Fig. 6d-e). We made the same analysis by considering the correlation for inside versus outside of cue-reward intervals (Fig. S6a,b). Note that all cue-reward intervals are enclosed in correct trial intervals.

To determine the number of variables contributing to the spiking of a given neuron, the prediction was performed for each of the possible 127 combinations of the seven variables (i.e. 7+21+35+35+21+1 from 1 to 7 variables). In the design matrix X, removed covariates were set as their mean over the entire session and the regressors β constructed from all seven covariates. We selected the combinations displaying the maximum correlation for each number of covariates and counted the number of times each covariate appeared in these best combinations. We then determined the minimum number of variables (or contributors) necessary to cross 90% of the largest value between the previously computed maximum correlations (Fig.6f).

To estimate which variables contributed the most to the spiking pattern, we looked at the relative loss in correlations after removing a single variable from the input to the model constructed from six covariates, and computed:

Relative loss in correlation = (correlation7covariates-correlation6covariates)/correlation7covariates. We computed the average relative loss per variable and per type of neurons (optogenetically identified DA neuron, putative DA neuron and non-cluster 5 neurons). On the other hand, we checked that the largest contributors really matter to the encoding. To do so, we compared the distributions of correlations when the encoder was given all 7 covariates to when the 3 strongest covariates were removed.

---

**Fig. S1.**
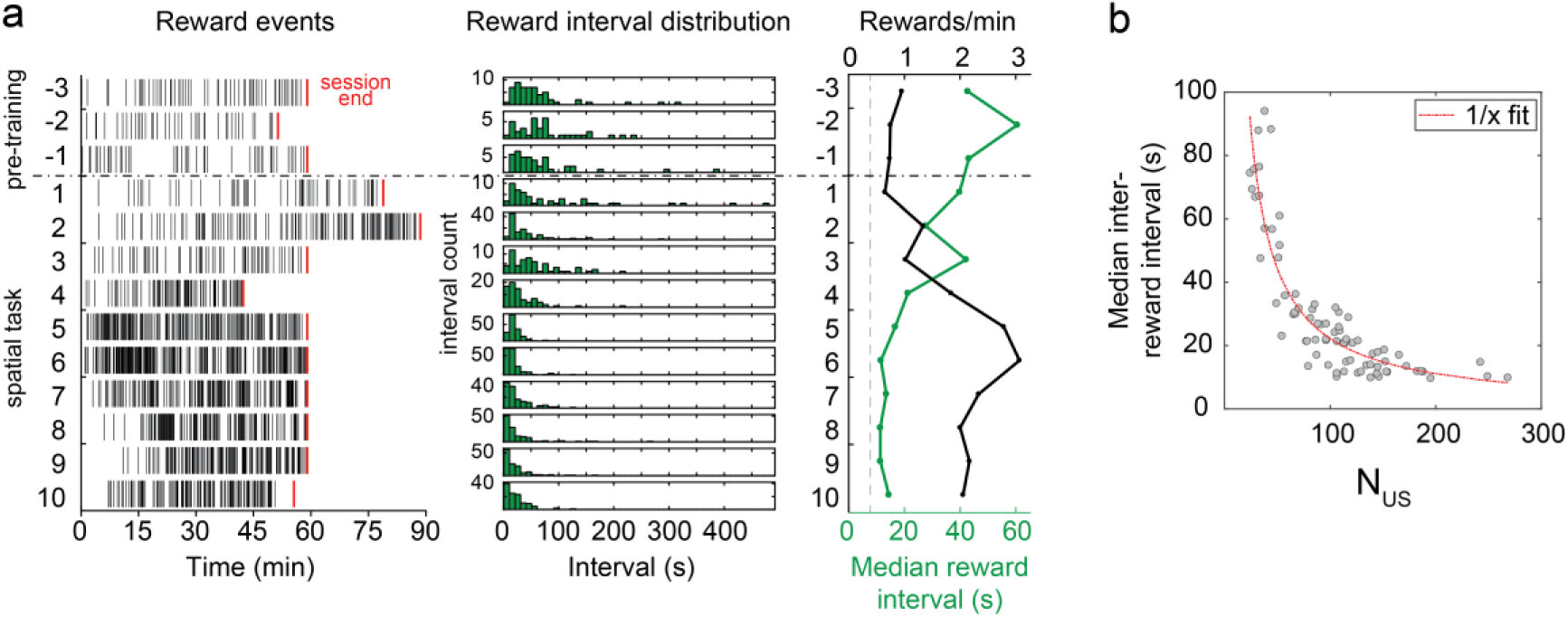
Median inter-reward interval as performance metric. **a,** Performance of one mouse across learning stages. Left: Temporal distribution of rewards for the last 3 pre-training sessions (sessions −3 to −1, top) and across 10 sessions of the spatial task (session 1 to 10). Red ticks mark the end of each session. Middle: Distribution of inter-reward time intervals across sessions, showing a gradual narrowing around short durations. Right: Median inter-reward interval (green) and average reward rate (black) across sessions. **b,** Median reward-interval plotted against the number of rewards obtained during the same session shows that a small median interval does reflect better performance and is not due to a low number of rewards. The red dashed line corresponds to a 1/x fit.

**Fig. S2.**
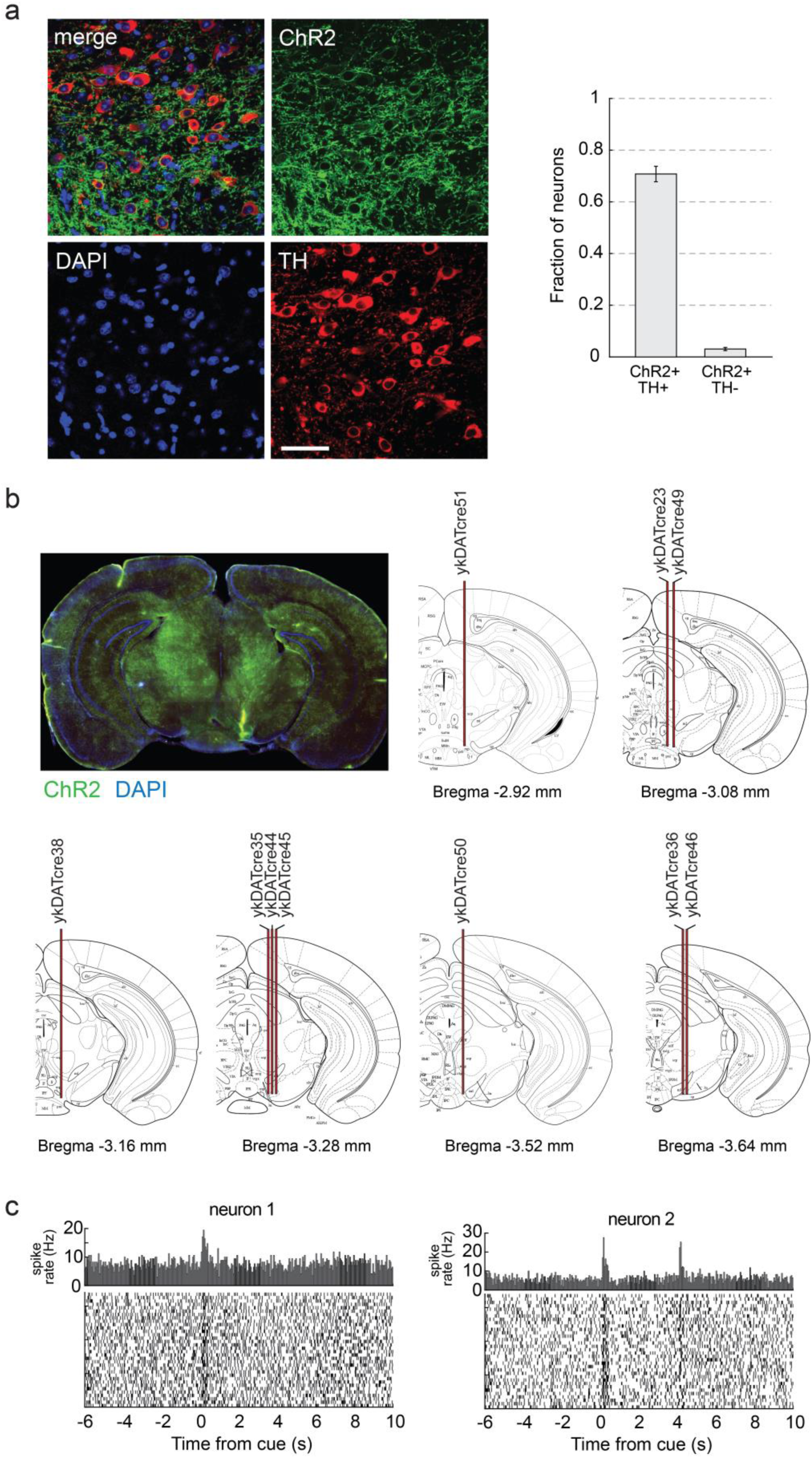
Post-mortem histology: ChR2 specificity & electrode tracks. **a,** Specificity of ChR2-expression compared to TH-staining (n=2 animals). Left: Example section of high-resolution confocal image of combined ChR2 expression (green), TH-(red) and DAPI staining (blue) (scale bar: 50µm). We found a high efficiency of transfection of TH-positive (TH+) neurons by ChR2-eYFP (70.8 % of ChR2+/TH+; 944 out of 1328 neurons) and a low proportion of leakage (3.1 % ChR2+/TH-; 43 out of 1328 neurons; bar graph shows mean ± s.e.m). **b,** Electrode tracks reconstructed after the end of the experiment. Top left: example of a slice showing ChR2-eYFP expression (green) and DAPI staining (blue) (scale bar: 2mm). The electrode track can easily be seen in the lateral part of the VTA. Right and below: individual electrode tracks from all animals traced on corresponding brain atlas slides ^78^. **c,** Two optogenetically identified DA neurons recorded during the same session of the spatial task. Top average spiking histogram and below raster plot of individual trials are both aligned to the cue. While the top neuron responded exclusively to the cue, the neuron below showed a larger response to the cue. The corresponding PSTHs are presented in the main Fig. 2d.

**Fig. S3.**
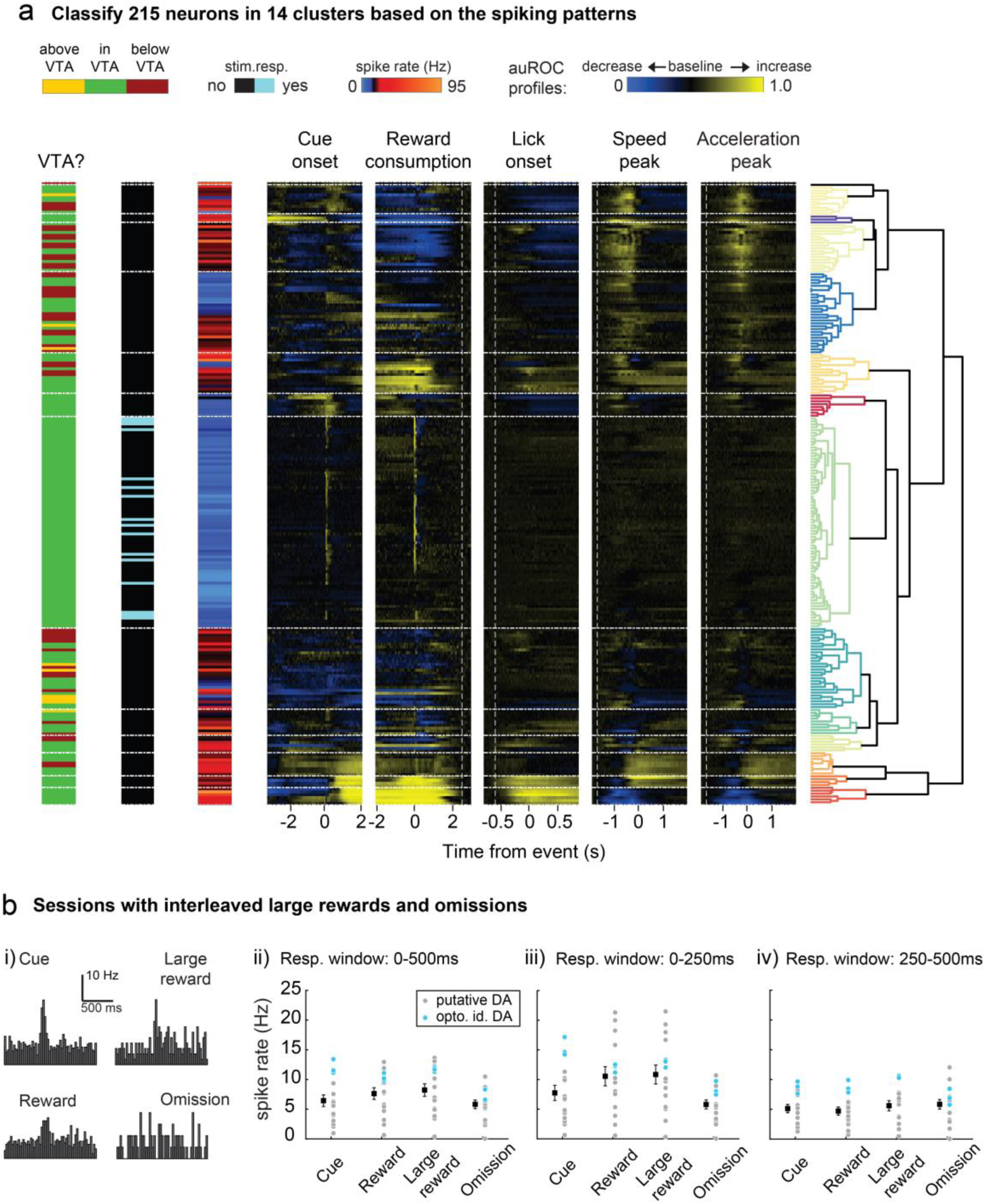
Pre-clustering analysis and impact of different reward types on the neural response. **a,** The hierarchical clustering applied to all 215 single-units produced 14 clusters among which the cluster #8 (n=73 neurons) contained all optogenetically identified DA neurons. Using this cluster, we looked in each mouse for recordings performed inside the VTA defined by the first and last occurrence of a DA neuron (yellow = above, green = inside and red = below the VTA). We used the resulting 159 neurons located in the VTA to perform a new hierarchical clustering yielding 9 clusters among which the cluster #5 contains the DA neurons (n=72; Fig. 3a). **b,** In a subset of experiments (n=3 animals), a small fraction of large rewards and omissions were randomly interleaved among correct trials. i) Average PSTHs for an example DA neuron to the cue and the 3 different outcomes (cue, large reward, omission). ii) Average responses for all DA neurons during a 500ms post-event window. For each condition, the mean ± s.e.m. across all neurons is represented by the black square. iii) Same representation of the early component of the post-event response (0-250ms; left) and iv) the late component of the post-event response (250-500ms; right) are plotted.

**Fig. S4.**
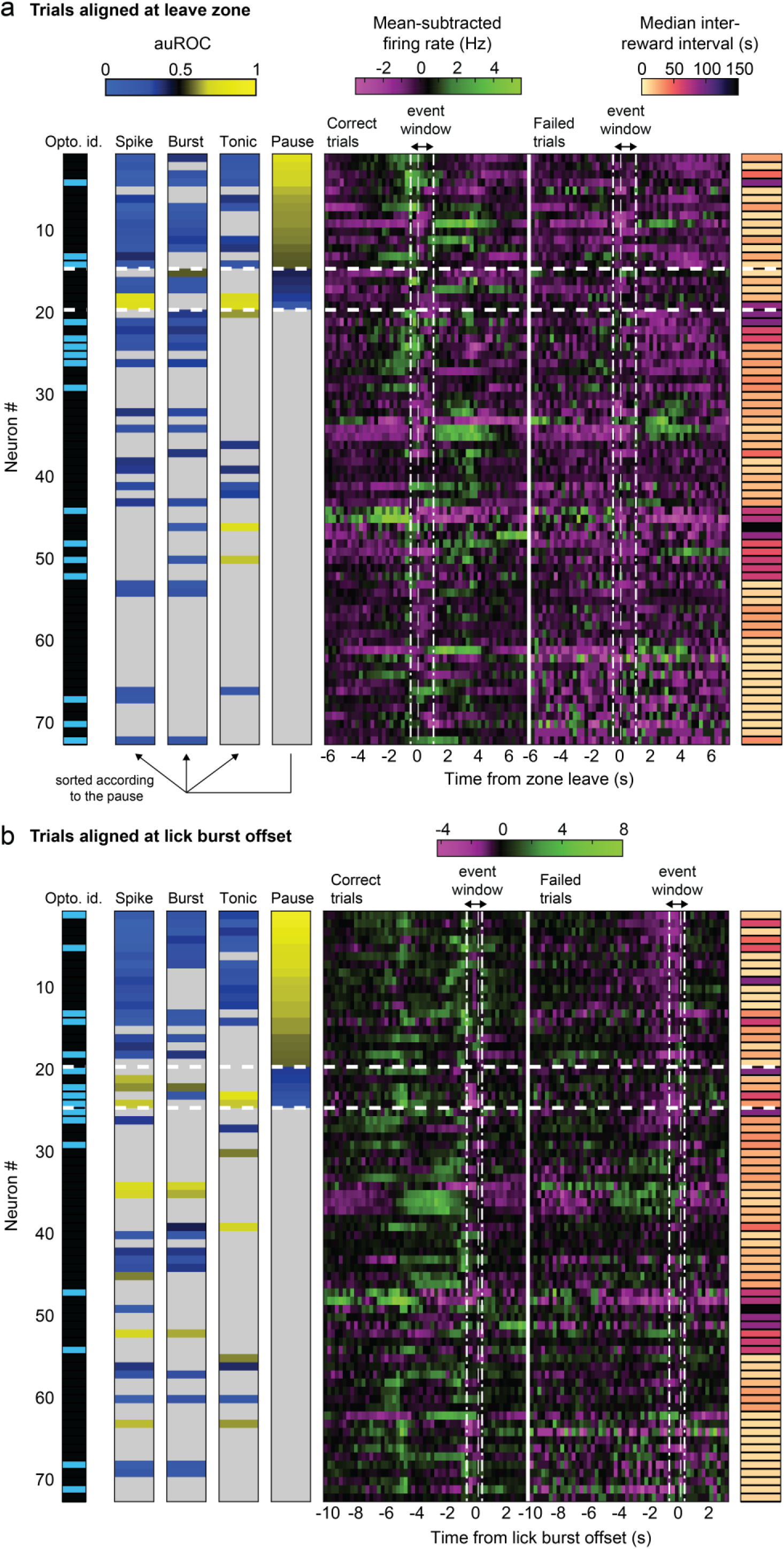
Neuronal activity for all optogenetically identified and putative DA neurons during correct and failed trials. **a,** Mean-subtracted firing rates at the time of the zone leave are shown for each of the putative and optogenetically identified DA neurons (cluster 5 neurons). For each neuron, optogenetic identification and the auROC analysis are indicated to the left. The performance of the animal during the corresponding recording session, represented as the median inter-reward interval, is plotted to the right. Neurons are sorted from top to bottom according to the auROC discrimination of the spike pauses. auROC values were quantified during the event windows for correct and failed trials highlighted by the vertical white dashed lines. We have 14/72 neurons with significant increase in firing pause and 5/72 neurons with significant decrease in firing pause. **b,** Same representation as in a, with an alignment of the neuronal activity and auROC analysis performed at the lick burst offset. As in a, neurons are sorted by the auROC value during firing pauses. We have 19/72 neurons with significant increase in firing pause and 5/72 neurons with significant decrease in firing pause. Thus, the lick burst offset is the event with largest number of pausing neurons among all four events. This is not surprising since at this time point mice know they failed the trial with certainty (i.e. they got no reward).

**Fig. S5.**
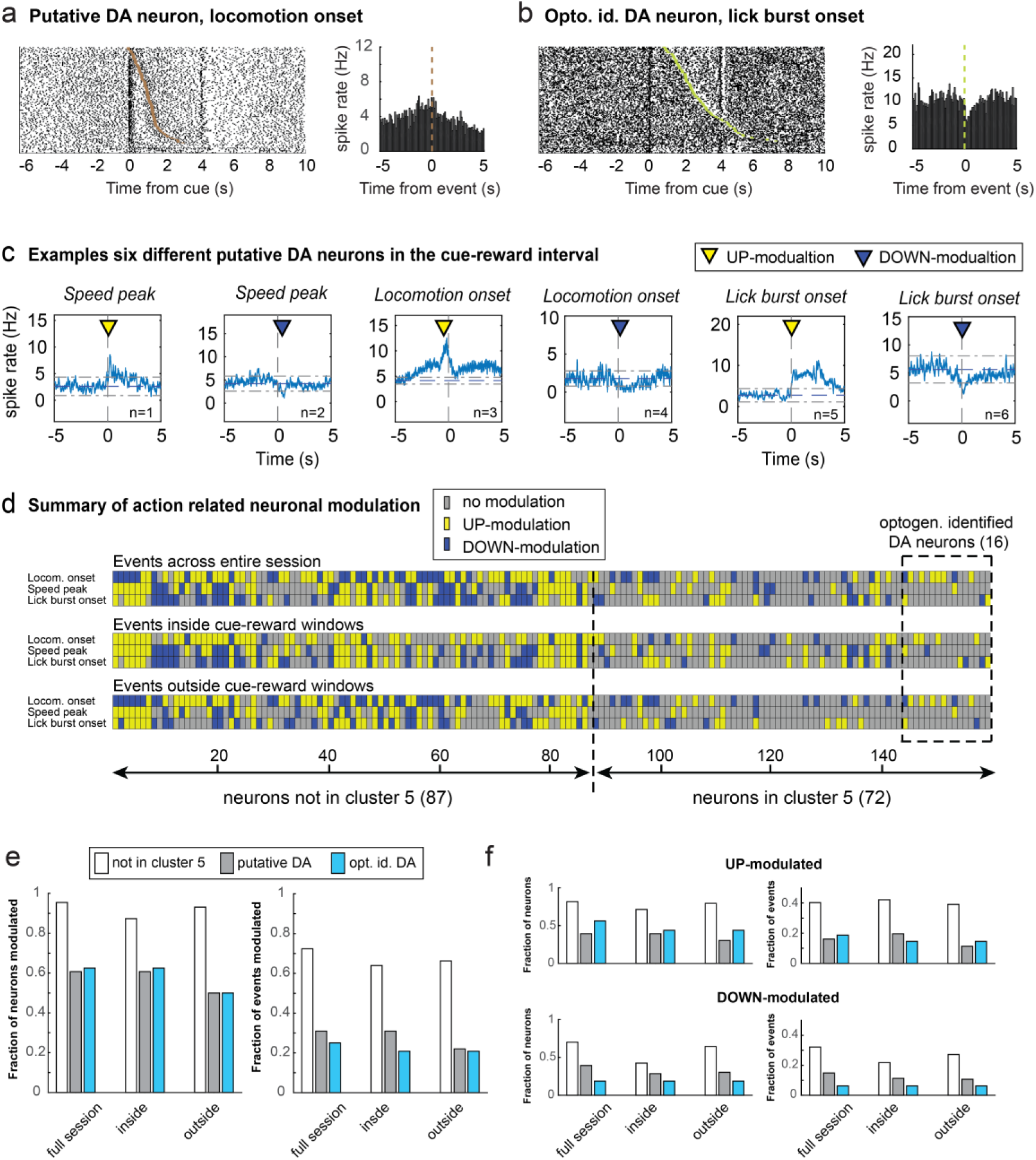
Neuronal modulation inside and outside of cue-reward intervals. a-b, Two action-modulated DA neurons showing clear phasic responses to cue and reward (the right DA neuron was optogenetically identified). While the left neuron (a) was transiently modulated by the locomotion onset, the second neuron (b) showed strong modulation to lick burst onset. The PSTHs are aligned to the onset of the motor output (bin size = 100 ms). Trials in the raster plot were sorted by increasing time interval between the cue and action onset. **c,** Example of six different putative DA neurons which are significantly modulated (either up: yellow arrow and down: blue arrow). **d,** Same representation as Fig.5c but excluding all motor events that are closer than 750 ms to the cue or reward. Detailed summary of spiking modulation by locomotion onset, speed peak and lick burst onset for all neurons grouped by cell-type (yellow: significant increase; blue: significant decrease; gray: non-significant). Top: For all events across the entire session. Middle: events between cue and reward. Bottom: events outside the cue-reward interval. **e,** Fraction of neurons (left) and fraction of events (right) modulated by motor parameters and grouped by cell-type for all events, between cue and reward and outside the cue-reward interval based on d panel. **f,** Fraction of neurons (left) and fraction of events (right) showing up-modulation (top) and down-modulation (bottom) by motor parameters and grouped by cell-type for all events, between cue and reward and outside the cue-reward interval (all values are available in Suppl. Table 1).

**Suppl. Table 1.**
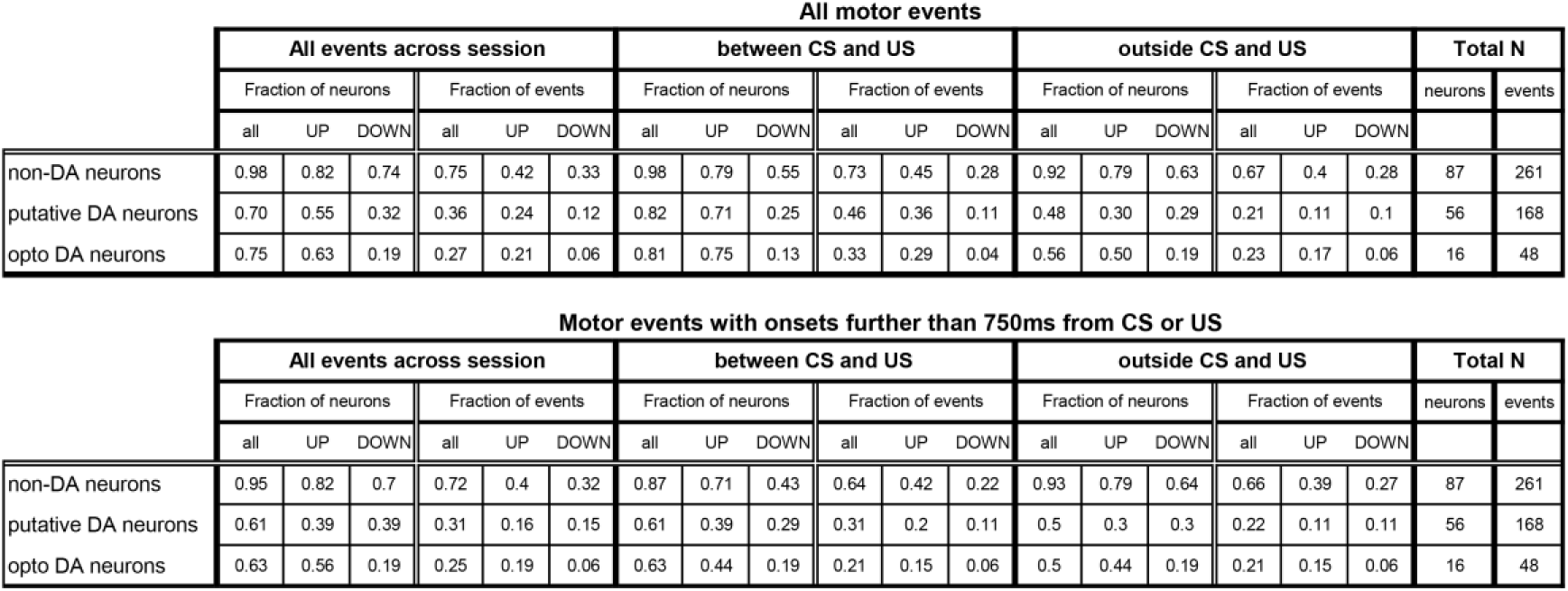
**, related to** Figure 5 **& S5: Analysis of movement related activation for all neurons.** All values used to plot panels in Fig. 5d & Fig. S5f-g. The fraction of neurons indicates how many cells have at least a single significant modulation to motor output. On the other hand. the fraction of UP-modulated neurons shows how many cells display at least one significant increase in spiking while the fraction of DOWN-modulated neurons shows the number of cell with at least one significant decrease in spiking. The fraction of events counts the number of all significantly modulated, UP-modulated or DOWN-modulated events grouped by cell-type (non-cluster 5 or non-DA neurons, putative DA or opto. Id. DA). The total number of events Nevents = 3 x Nneurons for each cell type.

**Fig. S6.**
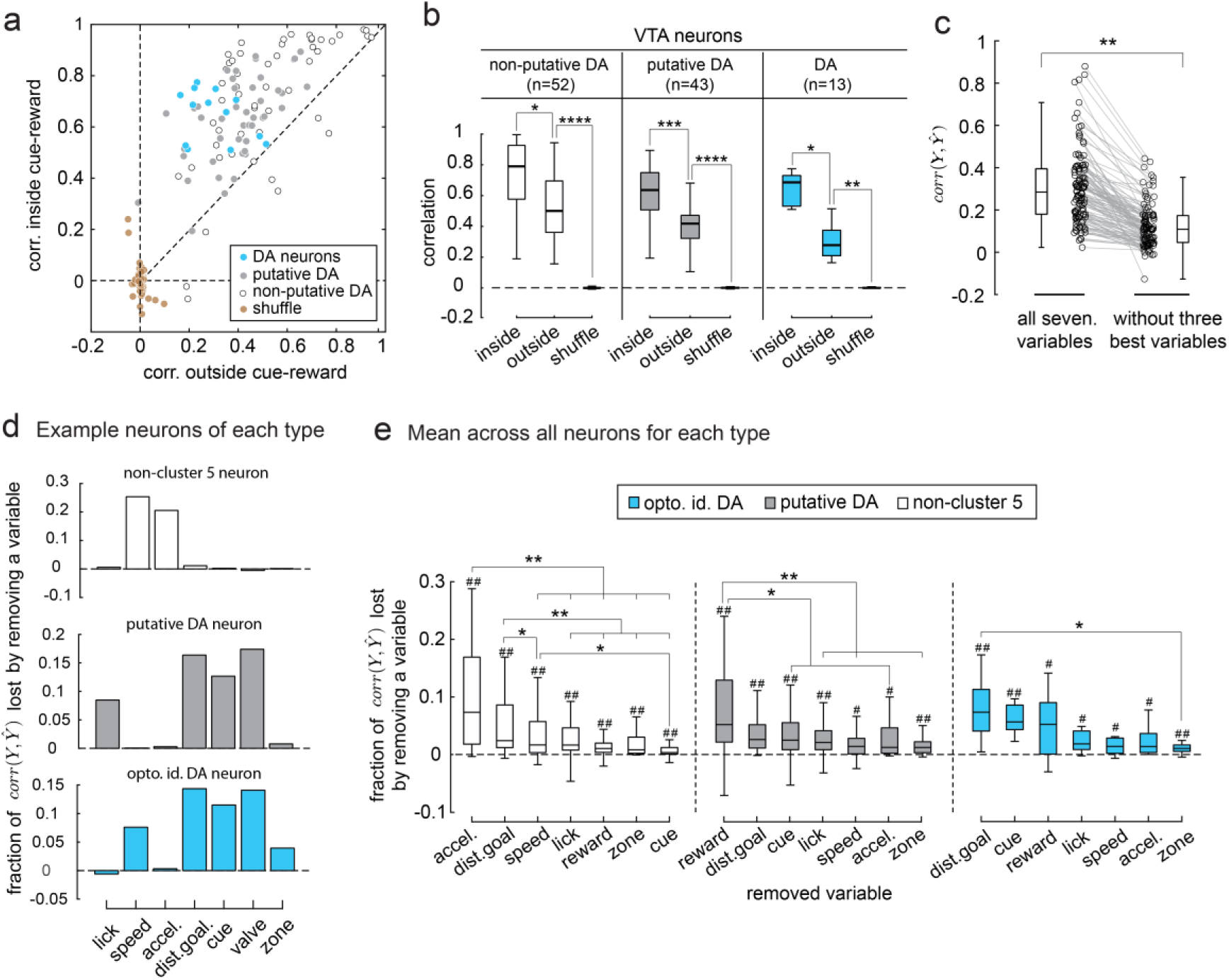
Compare encoding inside versus outside of cue-reward and estimate the relative importance of the different variables. **a,** Scatterplot comparing the correlation between the concatenated inside and outside pieces of the cue-reward interval (n=108). **b,** Using a Kruskal-Wallis nonparametric one-way ANOVA with post-hoc test Tukey-Kramer, we show that the variables encoding is stronger (and beyond the chance level) during the task execution (cue-reward interval) for all VTA neurons (*:p<0.05, **:p<0.01, ***:p<0.001, ****:p<0.0001). **c,** Comparing the distributions of correlations when the encoder was given all 7 covariates to when the 3 strongest covariates were removed by setting them to a constant value (the average of the covariate) showed a significant decrease (one-sided Kolmogorov-Smirnov test; p<<0.01), indicating that the predicted signal did encode these covariates. **d,** We estimate the relative importance of a given variable by the loss in correlation caused by removing it, i.e. we redo the encoding analysis but with covariate matrices construct from only six variables. The histograms show the relative loss in correlation for each variables (three example neurons, one of each type). The putative and opto. id. DA neurons display a similar profile apart that one better encodes speed (opto. id. DA) while the other encodes better licks (putative DA). On the other hand, the non-DA (or non-cluster 5) neuron only encodes two variables, namely speed and accelerations. This non-DA neuron does not encode the distance to goal showing that in this case it is not a confound for speed and acceleration. **e,** We compute the average loss of correlations across each type of neurons. We then sort the removed covariate (or variable) from the largest to the smallest contributions. The distance to the goal variable is of overall importance for all three groups of neurons. The average loss in correlation is significantly different from zero for all variables and neuronal type (t-test, #: p<0.05, # #: p<0.01). Note that the loss in correlation is not necessary significantly different between all variables (2-way ANOVA with Turkey-Kramer post-hoc test, *: p<0.05, **: p<0.01).

